# Exploring brain dynamics within the Approach-Avoidance Bias

**DOI:** 10.1101/2025.03.04.641451

**Authors:** Aitana Grasso-Cladera, Johannes Solzbacher, Debora Nolte, Peter König

**Affiliations:** Institut für Kognitionswissenschaft. Osnabrück University. Osnabrück, Germany; Department of Neurophysiology and Pathophysiology, Center of Experimental Medicine, University Medical Center Hamburg-Eppendorf, Hamburg, Germany

**Author notes:** **Corresponding Author**: Aitana Grasso-Cladera, Institute of Cognitive Science, University of Osnabrück. Wachsbleiche 27, 49090, Osnabrück, Germany. The authors contributed equally (Aitana Grasso-Cladera, Johannes Solzbacher).

**Keywords:** Approach-Avoidance Bias, Neural Dynamics, EEG, Approach-Avoidance Task, Event-Related Potentials, Frontal Alpha Asymmetry

## Abstract

Approach-avoidance behaviors (AAB) are fundamental mechanisms that guide interactions with the environment based on the emotional valence of stimuli. While previous research has extensively explored behavioral aspects of the AAB, the neural dynamics underlying these processes remain insufficiently understood. The present study employs electroencephalography (EEG) to systematically investigate the neural correlates of AAB in a non-clinical population, focusing on stimulus- and response-locked event-related potentials (ERPs). Forty-three participants performed a classic Approach-Avoidance Task (AAT) while EEG activity was recorded. Behavioral results confirmed the AAB effect, with faster reaction times in congruent compared to incongruent trials, as well for positive versus negative trials. ERP analyses revealed significant differences in the Valence factor, with early effects for stimulus-locked trials and late differences at the parietal-occipital region for response-locked trials. However, no significant effects were found for the Condition factor, suggesting that the neural mechanisms differentiating congruent and incongruent responses might not be optimally captured through EEG. Additionally, frontal alpha asymmetry (FAA) analyses showed no significant differences between conditions, aligning with the literature. These findings provide novel insights into the temporal and spatial characteristics of AAB-related neural activity, emphasizing the role of early visual processing and motor preparation in affect-driven decision-making. Future research should incorporate methodological approaches for assessing AAB in ecologically valid settings.

## Introduction

Approach-avoidance behaviors have been conceptualized as automatic responses generated by the organism regarding the subjective assessment of environmental components (e.g., objects, events, agents; Degner et al., 2021; McNaughton et al., 2016). Generally, the assessment is described in terms of the emotional valence of the cue (Bradley et al., 2001; Elliot, 2006; Lang, 1995; Renton et al., 2019), where positive or beneficial cues are more likely to be approached, and adverse or negative ones avoided (Aupperle et al., 2015; Eiler et al., 2019; Fridland & Wiers, 2018; Kenrick & Shiota, 2008). Empirical evidence supports the automaticity of these responses, demonstrating faster and more accurate reactions to stimuli congruent with the previously mentioned organism’s predispositions, which is known as the Approach-Avoidance Bias (AAB; Chen & Bargh, 1999; Czeszumski et al., 2021; Elliot, 2006; Krieglmeyer et al., 2013; Phaf et al., 2014; Zech et al., 2020). Moreover, the embodied nature of AAB underscores its evolutionary significance, with physiological mechanisms guiding these behaviors in realistic, dynamic contexts (Fridland & Wiers, 2018; Solzbacher et al., 2022). This interplay between emotional valence, automaticity, and embodiment highlights approach-avoidance behaviors’ evolutionary and adaptive role, offering valuable insights into how organisms navigate and respond to different environmental components.

Research into the AAB has largely prioritized behavioral dynamics, often focusing on reaction time differences as a key metric, while only a few studies have explored the neural mechanisms underlying these behaviors. Studies with humans and non-humans have determined structural and functional networks underlying approach and avoidance behaviors (Ironside et al., 2020; Lender et al., 2023; Wiers et al., 2014; Zorowitz et al., 2019). Within these networks, regions of the prefrontal cortex (PFC), specifically the anterior (aPFC) and adjacent ventrolateral areas, have shown greater activity while performing incongruent movements compared to congruent movements (Kaldewaij et al., 2017; Radke et al., 2015; Roelofs et al., 2009; Volman et al., 2011). This higher activity is linked to structural connections with the amygdala and its role in emotional processing and regulation (Andrewes & Jenkins, 2019; Sergerie et al., 2008), as well as the coordination of diverse cognitive processes (e.g., task-switching, control processing; (Ramnani & Owen, 2004). For the AAB, the PFC overrides automatic responses for incongruent reactions (Kaldewaij et al., 2017). The evidence highlights the pivotal role of the PFC in the cognitive processes underlying the AAB through its interplay with the amygdala.

Moreover, approach-avoidance behaviors can be understood as the result of the dynamic interplay between explicit and implicit information processing (Loijen et al., 2020; Strack & Deutsch, 2004a, 2004b). Explicit processing, such as under incongruent conditions in the Approach-Avoidance Task (AAT), involves deliberation and reflective reasoning. This type of processing is effortful, slower, and relies on cognitive control functions mediated by cortical frontal brain regions (Loijen et al., 2020). In contrast, implicit processing, evident in congruent AAT conditions, is driven by emotion-based information processing, influencing response timing and involving subcortical structures associated with emotional processing, such as the amygdala and ventral striatum (Loijen et al., 2020; Ochsner & Gross, 2005; Wager et al., 2008). According to Parsons et al. (2016) and Loijen et al. (2020), flexible and adaptive socio-emotional behaviors arise from the ongoing interaction between implicit and explicit processing mechanisms, including those underlying approach-avoidance behaviors. This framework raises questions about the temporal dynamics of these processes and their investigation through diverse methodological approaches, such as electroencephalography (EEG). Ultimately, approach-avoidance behaviors reflect a balance between explicit or emotion-driven and implicit or reflective mechanisms, where this interplay determines both the timing and nature of responses.

Although significant progress has been made in exploring the neural pathways associated with the AAB, inconsistent results in both temporal and spatial dynamics (Bamford et al., 2015; Lacey & Gable, 2021; Sege et al., 2024; Vecchio & Pascalis, 2020), have hindered a comprehensive understanding of the neural dynamics underlying evaluation, motivation, and decision-making processes. Furthermore, many articles exploring brain activity have focused on clinical populations (Ellis et al., 2019; Fechtner, 2013; Sege et al., 2024; Vecchio & Pascalis, 2020); however, neural dynamics within the AAB on healthy subjects are not fully understood. This exploratory article uses EEG to systematically investigate neural dynamics in healthy subjects while performing a classic Approach Avoidance Task (AAT). To this end, we partially replicated the experimental paradigm implemented by Solzbacher and colleagues (2022) and collected EEG data while participants performed the task. To our knowledge, this is the first study to explore neural differences associated with the AAB using both stimulus- and response-locked trials in a non-clinical population. By systematically investigating differences between stimulus- and response-locked ERPs, we aim to deepen our understanding of human behavior by providing insights into the timing and nature of cognitive and emotional processes underlying the AAB.

## Methods

### Participants

A total of 43 participants (26 males) aged 19 to 36 years (mean age = 25.30, SD = 3.57) completed the experiment. All participants were right-handed, had normal or corrected-to-normal vision, and were advanced or native speakers of English. Before the experiment, written informed consent was obtained from all participants. They received course credit as compensation for their participation. All instructions were provided and explained in English. The study was approved by the Ethics Committee of Osnabrück University.

### Stimuli

The stimulus set included 87 full-color pictures belonging to the International Affective Picture System (Lang et al., 1997), chosen to ensure comparability and consistency with prior research (Czeszumski et al., 2021; Kaspar et al., 2015; Solzbacher et al., 2022). The images were selected considering the cumulative evidence and their reliability for evoking emotional states across diverse populations and contexts (Branco et al., 2023; Lang et al., 1993, 1998; McManis et al., 2001) and based on ratings of their *Valence* (pleasant vs unpleasant) from the Self-Assessment Manikin (SAM) scale. Of the 87 pictures, 44 had valence ratings below three points (unpleasant stimuli)^1^; the other 43 had valence ratings above seven (pleasant stimuli)^2^. The pictures featured diverse content and were displayed at their original resolution (1024 x 768 pixels), centered against a gray background (RGB values: 182/182/182).

### Apparatus

The stimuli were presented on a 24-inch LCD monitor (BenQ XL2420T; BenQ, Taipei, Taiwan) positioned 80 cm in front of the participants. The display was set to a resolution of 1920 × 1080 pixels with a refresh rate of 114 Hz. The experiment was implemented using Psychtoolbox V3 (Kleiner et al., 2007) in Matlab R2016b (MathWorks). Behavioral data were collected through a Logitech Extreme 3D Pro joystick, selected for its capacity to facilitate body-related gestures. Participants used the joystick to simulate approach-avoidance behaviors by pushing and pulling, allowing the setup to examine embodied elements of the automatic approach-avoidance bias (Solzbacher et al., 2022). Consistent with previous studies, reaction times were measured from the onset of the joystick’s push or pull motion (Solarz, 1960; Solzbacher et al., 2022).

Brain activity was collected using EEG. A 64-channel Ag/AgCl electrode system with a Waveguard cap (ANT, Netherlands) and a Refa8 amplifier (TMSi, Netherlands), managed through the asa-lab acquisition server (v4.9.4) on a Dell laptop (Dell Inc.; Windows 7, 32-bit; Intel(R) Core(TM) i5-3320M CPU). Data were sampled at 1024 Hz and recorded using common-reference. A ground electrode was positioned under the left collarbone. EOG electrodes were positioned above and under the left eye. Impedances were kept below 10 kΩ. Triggers for stimulus presentation and movement onset were sent using a parallel port.

### Procedure and Design

The experimental design partially replicates a previous study by Solzbacher and colleagues (2022). Participants were seated in front of a computer screen and presented with a series of pictures displayed one at a time. From the different variations of the approach-avoidance tasks described in the literature, we opted to implement explicit instructions (i.e., active evaluation of the valence of the stimulus) since they appear to yield the most consistent results (Phaf et al., 2014; Van Dessel et al., 2016). Hence, participants were instructed to classify the pictures as either positive or negative. This classification was made by performing a joystick movement either toward their body (i.e., a pull movement) or away from their body (i.e., a push movement). All participants performed the task with their dominant hand and were asked to respond to each picture as quickly and precisely as possible. It was not possible to rectify and correct response mistakes, and no feedback was delivered.

The task consisted of two blocks with different instructions (congruent or incongruent). The order of the blocks was randomized for each participant. In the congruent block, participants were instructed to react to positive pictures by performing a *pull* movement towards their body (approach) and to react to negative pictures with a *push* movement away from their body (avoid). In the incongruent block, the participants were instructed to approach the negative images and avoid the positive pictures. To ensure participants remained aware of the current block’s instructions, reminders were displayed every 20 trials.

Each trial began with a fixation cross displayed at the center of the screen for 3 seconds (±0.5 seconds). Then, the picture appeared and remained on the screen until the participant moved using the joystick. At the end of each trial, there was a pause of 0.5 seconds. Participants were presented with 40 images of different emotional valence, equally distributed, in each block. At the beginning of each block, the first four images were test trials and were excluded from the analysis. Each image was presented once during the entire experiment. The order of the stimuli was randomized between blocks and subjects.

Following previous studies (Czeszumski et al., 2021; Solzbacher et al., 2022), a “zoom effect” was incorporated to augment the impression of an approach or avoidance behavior. The “zoom effect” consists of dynamically altering the image sizes in response to joystick movements—images increased in size during approach (pull) movements and decreased in size during avoidance (push) movements. This feature started when the participants initialized the movement of the joystick and continued enlarging or reducing the image’s size at a fixed speed. The “zoom effect” was programmed in MATLAB’s Psychtoolbox V3 (r2017a; MathWorks Company; adjusted from Czeszumski et al., 2021). Figure 1 provides a graphical representation of the task.

**Figure 1.**
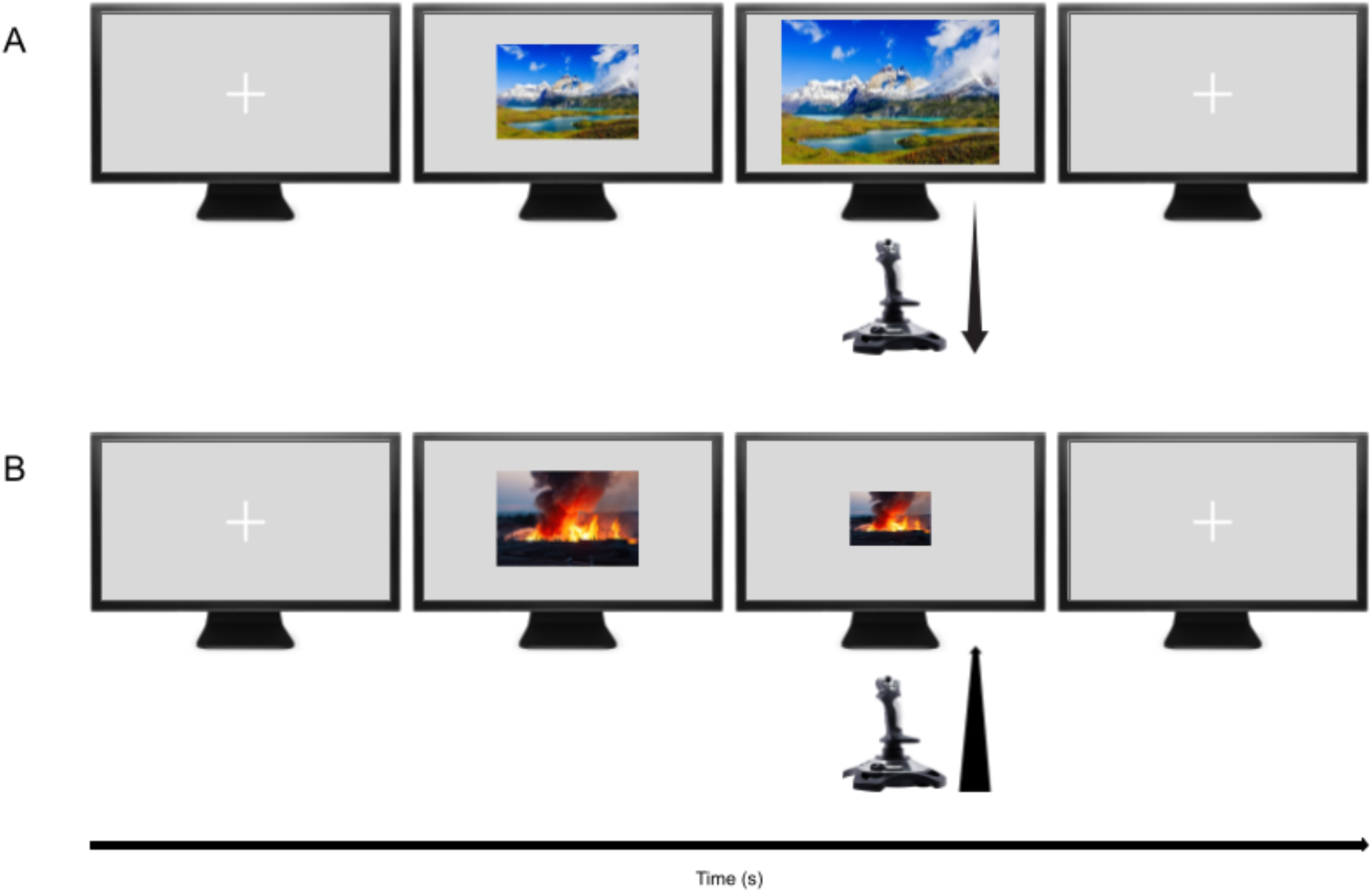
Graphical representation of the experimental paradigm^3^. Panel A and B show an example of a congruent block, where the participant performs a *pulling* movement with the joystick on a positive picture, which zooms in (A) or performs a *pushing* movement on a negative picture, which zooms out (B).

Before the picture trials, participants performed 20 no-stimulus trials in which they were instructed to push or pull the joystick whenever the fixation cross changed color. This was done to obtain baseline data while using the joystick and to help participants familiarize themselves with the movement.

### Preprocessing

#### Behavioral Data

The collected data was divided into two conditions: congruent and incongruent. We visually inspected the reaction time data, indicating a positive skew (i.e., right-skewed distribution) for both conditions. Such a distribution shape is characteristic of reaction-time experiments, where a concentration of faster responses and a tail of slower responses are commonly observed (Morís Fernández & Vadillo, 2020; Rousselet & Wilcox, 2018). To address outliers, we first implemented a filtering procedure. Since the minimum reaction time for processing visual stimuli is typically around 200 ms (Jain et al., 2015; Otaki & Shibata, 2019), we identified all trials with reaction times below 150 ms as unintentional responses, excluding 16 trials across participants (4 in the congruent condition and 12 in the incongruent condition). These rapid responses were likely unrelated to the task and stimulus processing, potentially arising from factors such as the influence of hand position on the joystick, where resting hand weight may have hindered a return to the neutral position. Furthermore, in some trials, participants exhibited reaction times exceeding two standard deviations from the mean, resulting in a longer “tail” distribution. We applied a 2-standard-deviation Winsorizing process to adjust the dataset’s distribution while preserving valuable information. The 2-standard-deviation adjustment affected 112 trials across participants, representing 3.38% of the entire dataset (55 trials in congruent and 57 in incongruent conditions). Finally, participants with performance lower than 90% accuracy (N = 2) were excluded from further behavioral and EEG analyses.

#### EEG Data

EEG data preprocessing was conducted in the MATLAB (R2024b) environment using EEGLAB (Delorme & Makeig, 2004; version 2024.1) with a custom script adjusted from Schmidt and Nolte (2024). First, the data was imported into MATLAB, ensuring double precision for all preprocessing steps (Bigdely-Shamlo et al., 2015). Non-empirical segments (e.g., pre-task intervals) were removed. Channel labels were standardized to the 10-5 BESA system, and channels with no recorded data were excluded.

We applied a Hamming window (Widmann et al., 2015) and filtered the data with a low-pass filter at 128 Hz and a high-pass filter at 0.5 Hz using the *pop_eegfiltnew* function. Subsequently, the data was downsampled from 1024 Hz to 500 Hz, after which we applied the *Zapline Plus* plugin (de Cheveigné, 2020; Klug & Kloosterman, 2022) to attenuate line noise, specifically addressing spectral peaks around 50 Hz. Data was then re-referenced to the average to establish a consistent baseline.

The *clean_rawdata* function was used to automate cleaning of noisy channels (mean = 3.674, std = 2.53, min = 0; max = 9) and segments (mean = 5.413, std = 6.57; min = 1, max = 32) (Kothe, 2014). Given the higher-than-expected noise level, we applied a conservative burst criterion of 20, representing the standard deviation threshold for burst removal via Artifact Subspace Reconstruction (ASR). Following noisy channel removal, data was again re-referenced to the average.

Independent component analysis (ICA) was conducted using the Adaptive Mixture of Independent Component Analyzers (AMICA) plugin (version 15; Palmer et al., 2012) to isolate and remove components associated with muscle, ocular, cardiac, and residual line or channel noise. For ICA preprocessing, a temporary high-pass filter at 2 Hz was applied to improve component estimation (Dimigen, 2020). Independent components (ICs) with greater than 80% muscle-related activity or over 90% other noise, as identified by ICLabel (Pion-Tonachini et al., 2019), were automatically rejected (mean = 7.372, std = 4.14, min = 2, max = 18). Then, EEG sensor-level data was reconstructed from the ICs and robustly identified as brain-related. Missing channels were interpolated using spherical interpolation. This procedure was implemented consistently across all subjects, ensuring uniform preprocessing before statistical analysis. Participants with less than 90% valid trials (N = 1 for stimulus-onset trials and N = 2 for movement-onset trials) were removed from further analyses.

Finally, we implemented a linear model using the Unfold toolbox (Ehinger & Dimigen, 2018) to correct for the effect of overlapping events due to the experimental design. In the model, we use both picture onset and movement onset as event factors, with the levels of the 2×2 design (i.e., congruent-positive, congruent-negative, incongruent-positive, incongruent-negative). The overlap correction adjustment was applied for -800 to 1000 ms surrounding picture onset and -800 to 800 ms surrounding movement onset.

### Analyses

#### Behavioral Analysis

We employed a linear mixed model (LMM) to analyze the behavioral data, which assumes that the data follow an underlying normal distribution. Since our reaction time data exhibits an ex-Gaussian distribution (Park & Hyun, 2014), we applied a base-10 logarithmic transformation (log₁₀) to the data to meet the normality assumption. We performed all subsequent behavioral analyses on the log-transformed reaction time data. The LMM was implemented in MATLAB using the *fitglme* function, with Maximum Pseudo-Likelihood (MPL) as the method fit. We assumed constant degrees of freedom for the model. Effect sizes were computed using Cohen’s d (Cohen, 1988; Durlak, 2009; Goulet-Pelletier & Cousineau, 2018), calculated by dividing the coefficient of each prediction by the corresponding standard error. We model reaction times by the condition (i.e., congruent/incongruent) and the emotional valence of the picture (i.e., pleasurable/unpleasurable) factors, which were defined as fixed effects. We tested the interactions between the mentioned factors. All predictors were included using the effect coding scheme (Te Grotenhuis et al., 2017). Furthermore, we included the grouping variables *subject* and *picture* as random effects to account for variability arising from individual participant differences and picture-specific factors not explained by the valence rating. Equation 1 presents the Wilkinson notation of the implemented model.

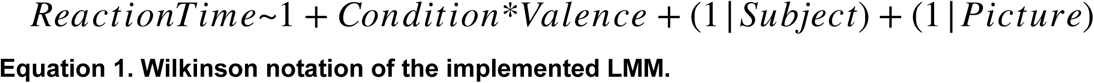

#### EEG Analysis: Time Series Domain

To explore differences across conditions at all electrodes and time points, we performed a two-factor repeated measures ANOVA (2×2: Condition x Valence), with a significance level set at .05. To address the multiple comparison problems, we implemented a cluster-based permutation test using threshold-free cluster enhancement (TFCE) as described on the ept_TFCE toolbox (Mensen & Khatami, 2013). We performed 10.000 permutations, randomizing the data across all factors and levels for each permutation, and then applied a two-factor repeated measures ANOVA. Subsequently, the resulting F-values were enhanced using TFCE with parameters E = .666 and H = 1, following recommendations for F-statistics (Mensen & Khatami, 2013). This process generated an empirical null distribution of TFCE-enhanced F-values, derived by recording the maximum F-value across all channels and time points for each permutation. Observed TFCE-enhanced F-values were then evaluated against this null distribution, with statistical significance identified as values exceeding the 95th percentile of the null distribution.

#### EEG Analysis: Frontal Alpha Asymmetry (FAA)

The EEG signal’s power spectral density (PSD) was computed using Welch’s method to estimate the signal power distribution across frequencies. The activity post-picture onset of each epoch was analyzed individually by applying a Hamming window of 1-second duration (500 samples) with 50% overlap to mitigate spectral leakage. The PSD for each epoch was calculated using a 1024-point Fast Fourier Transform (FFT), resulting in a frequency resolution of approximately 0.49 Hz. The PSDs were averaged across all trials to obtain an average spectrum. Alpha band power (8–12 Hz) was extracted by integrating the PSD over the specified frequency range. This approximation provides a robust estimate of the spectral content while accounting for inter-epoch variability. Then, we calculated Frontal Alpha Asymmetry by subtracting the frontal activity on the alpha band of the right and left hemispheres (Fox et al., 1995; Lacey & Gable, 2021). We computed FAA over single electrodes and left-right clusters. Finally, we implemented non-parametric statistics (Wilcoxon Signed-Rank test) to evaluate differences between pleasant and unpleasant trials.

## Results

### Behavioral Data

First, we investigated whether the collected behavioral data reproduces previously observed patterns of reaction time data as a function of condition and valence. For this, we applied a Linear Mixed Model. Interestingly, the results from the model revealed a significant main effect of *Condition*, t(3192) = -5.206, p < .0001. Specifically, the log-transformed reaction time data for the congruent condition was estimated to be lower than for the incongruent condition, with a coefficient of −.008 (95% CI: [-.0111, .0050]). This corresponds to the congruent condition having reaction times of approximately .9817 compared to those in the incongruent condition, derived from the exponential of the log-transformed coefficient (Figure 2). This indicates that participants responded significantly faster in the congruent condition than the incongruent condition, with a difference of approximately 1.83%. The effect size for the *Condition* factor was d = -5.2 (SE = .0015), corresponding to a large effect size (Cohen, 1988). Additionally, we found a significant main effect of *Valence* t(3192) = -2.95, p = .0032, which indicated that the log-transformed reaction time data for pleasant pictures was estimated to be lower than for unpleasant pictures, with a coefficient of −.0118 (95% CI: [-.0196, -.0039]). This corresponds to the condition of the pleasant picture having reaction times of approximately .9732 than those in the condition of the unpleasant picture, derived from the exponential of the log-transformed coefficient (Figure 2). Hence, participants performed a movement significantly faster for pictures rated as pleasant when compared to the unpleasant ones (2.68%; Figure 2). For the *Valence* factor, the effect size was d = -2.95 (SE = .0040), corresponding to a large effect size (Cohen, 1988). The results from the tested interaction of the relevant factors (*Condition* * *Valence*) showed no significant differences (p = .0652). Finally, the Interclass Correlation Coefficient was .4103 for the grouping variable *Participants* and .0825 for *Pictures*. Overall, the behavioral data results align with the AAB reaction time pattern.

**Figure 2.**
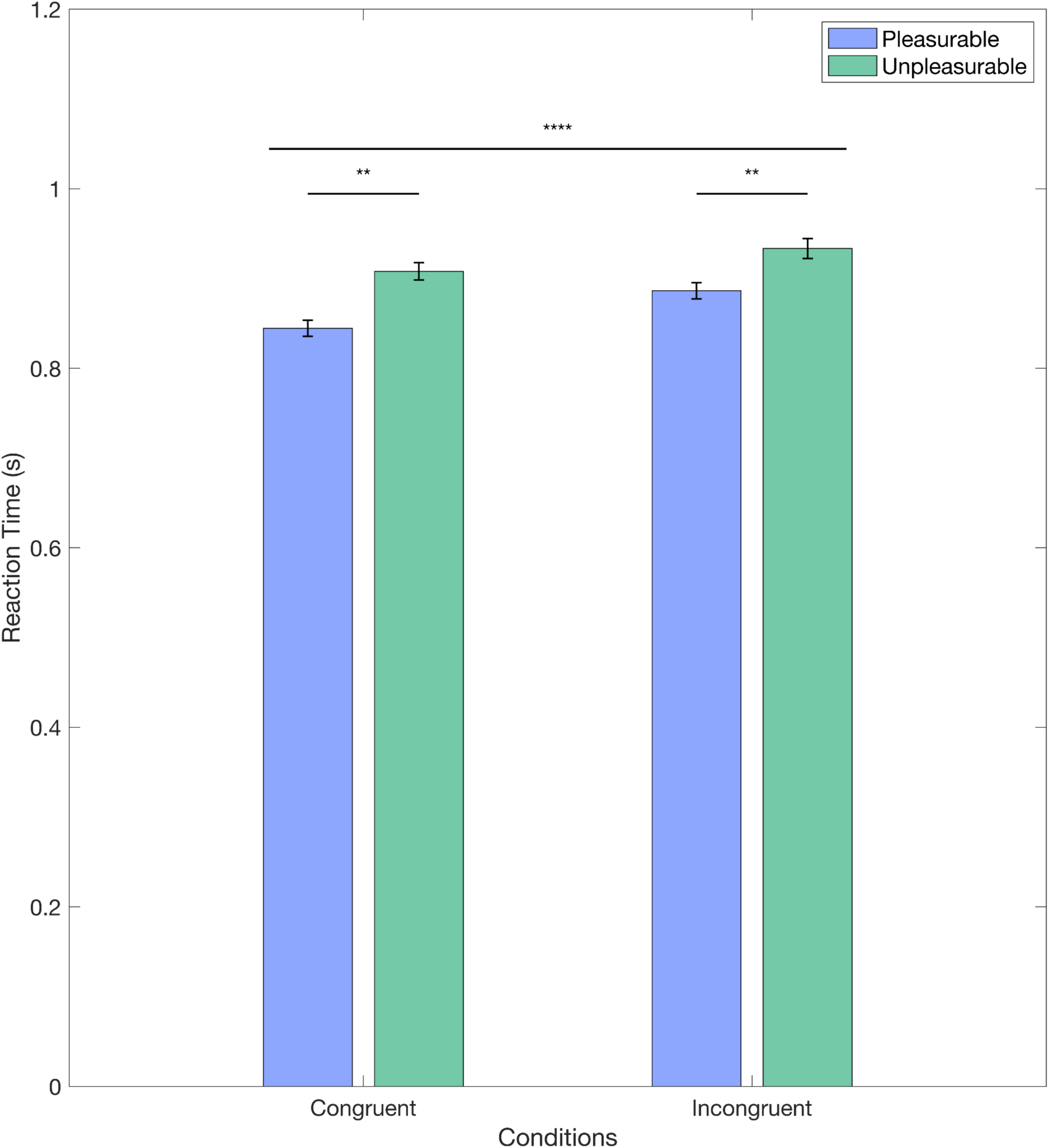
Main Effects for the LMM for Behavioral Data. The figure displays the mean reaction times for each condition and the standard error for the non-log transformed data. Significant differences for the *Condition* (congruent vs incongruent) and *Valence* (positive vs negative) factors are displayed here.

### EEG Analysis: Time Series Domain

Regarding the time-series domain, we aimed to explore differences across conditions at all electrodes and time points for stimulus- and movement-onset ERPs. For this, we first preprocessed the data by implementing a linear model using the Unfold toolbox (Ehinger & Dimigen, 2018) to correct for the effect of overlapping events (stimulus and movement onset) due to the experimental design and then proceeded to epoch the data (-800 to 1000 ms for stimulus-onset ERPs and -800 to 800 ms for movement-onset ERPs). A total of 3060 trials were considered valid for stimulus-onset and 2987 for movement-onset. Considering the large number of electrodes and time points, we conducted a mass univariate analysis (Rousselet & Pernet, 2011; Stropahl & Debener, 2017; Winward et al., 2022) to explore condition-specific differences along electrodes and time. We used threshold-free cluster enhancement (TFCE; Mensen & Khatami, 2013) as a correction for multiple comparisons with a significance level at alpha < .05. Following this procedure, we were able to perform data-driven analyses of complex EEG data on the time-series domain.

We applied the described procedures for investigating different conditions (Congruent-Positive, Congruent-Negative, Incongruent-Positive, Incongruent-Negative) for stimulus- and movement-locked ERPs. We found significant differences in the *Valence* factor and the interaction of factors for both stimulus- and response-locked ERPs, and there was no significant cluster for the *condition* factor. For stimulus-locked analyses, we found the peak for the significant cluster for the *Valence* factor at 86 ms after stimulus onset at channel M2, while for the interaction of factors, the peak was at 582 ms after stimulus onset at channel P6. For response-locked ERPs, we found the peak of the cluster for the *Valence* factor at 2 ms after movement onset at channel P5 and 108 ms post movement onset at channel POz for the interaction of factors. Figure 3 displays each significant cluster peak’s ERPs. The results from stimulus- and response-locked ERPs show significant clusters for the *Valence* factor and the interaction of factors but not for the *Condition* factor.

**Figure 3.**
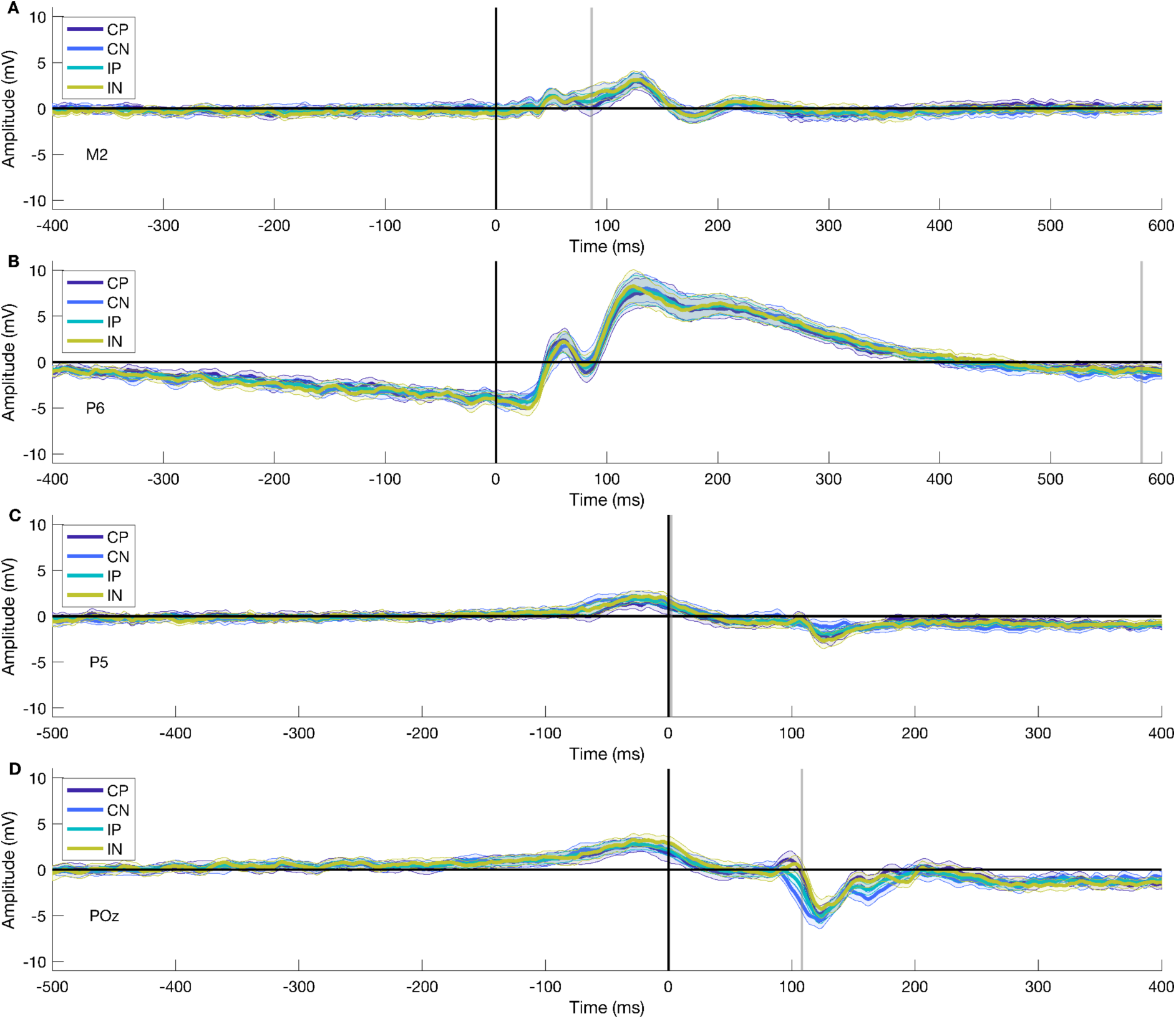
ERPs for all peaks of significant clusters. Panels A and B represent stimulus-onset ERPs for *Valence* and *Interaction,* respectively. Panels C and D display movement onset ERPs for *Valence* and *Interaction,* respectively. The gray line represents the temporal peak of the cluster. For visualization purposes, we have shortened the temporal window to -400 to 600 ms for picture onset trials and -500 to 400 ms for movement onset trials.

The spatial distribution of the cluster’s peak varied for every ERP type and significant level. For stimulus-locked ERPs in the *Valence* factor, the peak of the cluster encompassed 36 electrodes distributed across frontal, central, and posterior areas (Figure 4A), while for the interaction of factors, the peak of the cluster included 10 electrodes with a parietal-occipital distribution (Figure 4B). Furthermore, for response-locked ERPs, the extent of the peak of the significant cluster was 7 electrodes, with a parietal-occipital distribution (Figure 4C), and for the interaction of factors, there were 50 electrodes included in the peak of the cluster distributed across different topographic areas (Figure 4D). Overall, the topographical extension of the significant clusters varied for stimulus- and movement-locked ERPs and for the significant levels.

**Figure 4.**
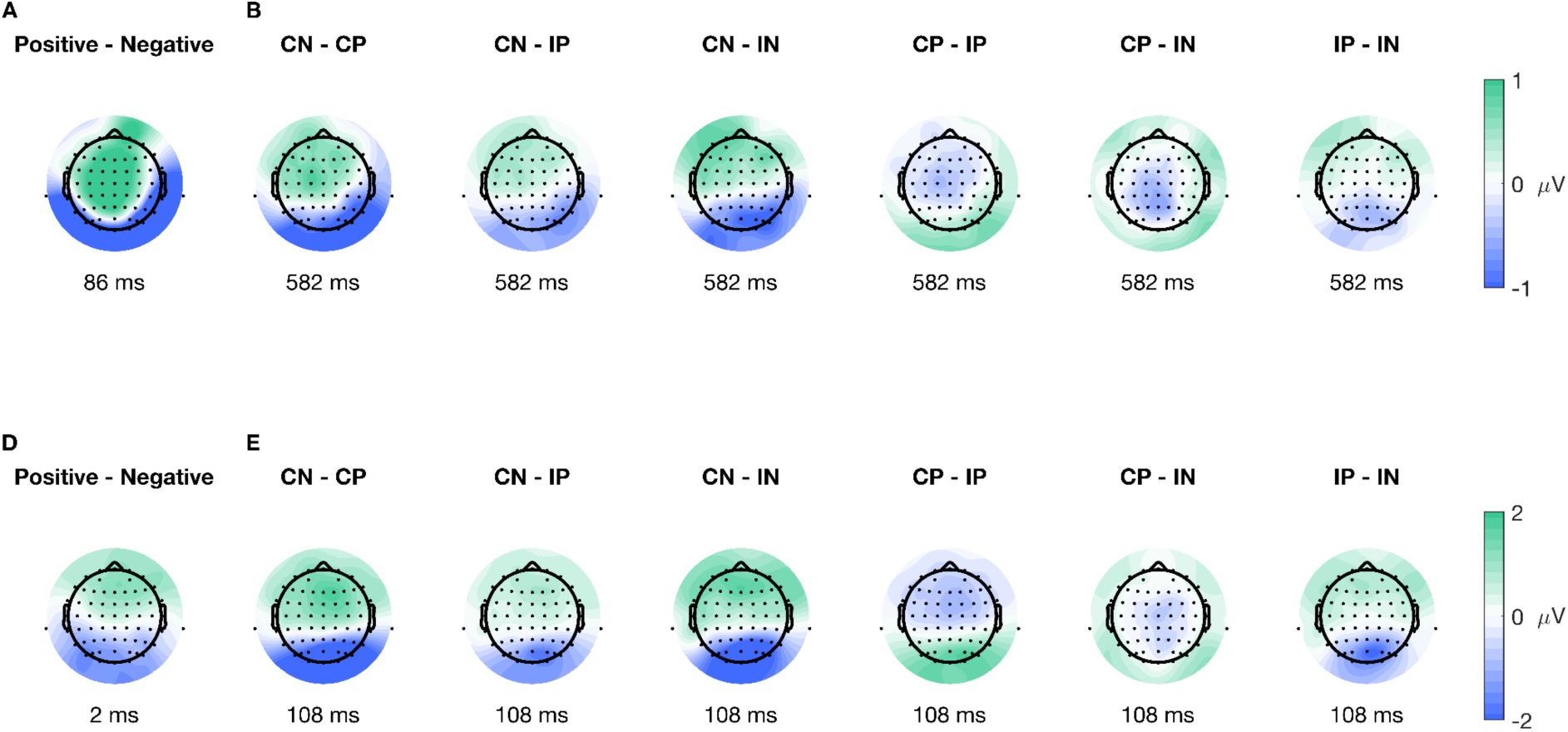
Topographic plots for all peaks of significant clusters. Panels A and B illustrate the average difference between conditions for the cluster peak on stimulus-onset ERPs, A for the *Valence* factor, and B for the interaction. Similarly, panels C and D show the average difference between conditions for the cluster peak on movement-onset ERPs, with C displaying the *Valence* factor and D the interaction.

### Stimulus-Locked Cluster Characteristics

In the following sections, we will describe the results, specifically the cluster extent, for the *Valence* factor and the interaction of factors for ERPs aligned to picture-onset, given that there are no differences for the *Condition* factor.

The mass univariate analysis showed differences in the *Valence* factor of the stimulus-onset ERPs. We performed a test for significant differences using TFCE (2×2 ANOVA, alpha < .05), which revealed a significant cluster highly compatible with an effect spanning from 30 to 126 ms post-picture onset (Figure 5A-D), encompassing a total of 57^4^ out of the 64 electrodes. The peak of the cluster was found at channel M2, at 86 ms after picture onset (Figure 5A and the second difference plot in 5E), presenting a median difference between positive and negative trials of -1.921 µV (std = 1.624, [-5.703 1.181]) of this time-electrode combination. The peak of this cluster is congruent with the P100 component, associated with early visual preprocessing. This result shows significant early differences after picture onset regarding the emotional valence of the stimuli; the difference is driven by higher amplitudes observed during negative trials than positive ones.

**Figure 5.**
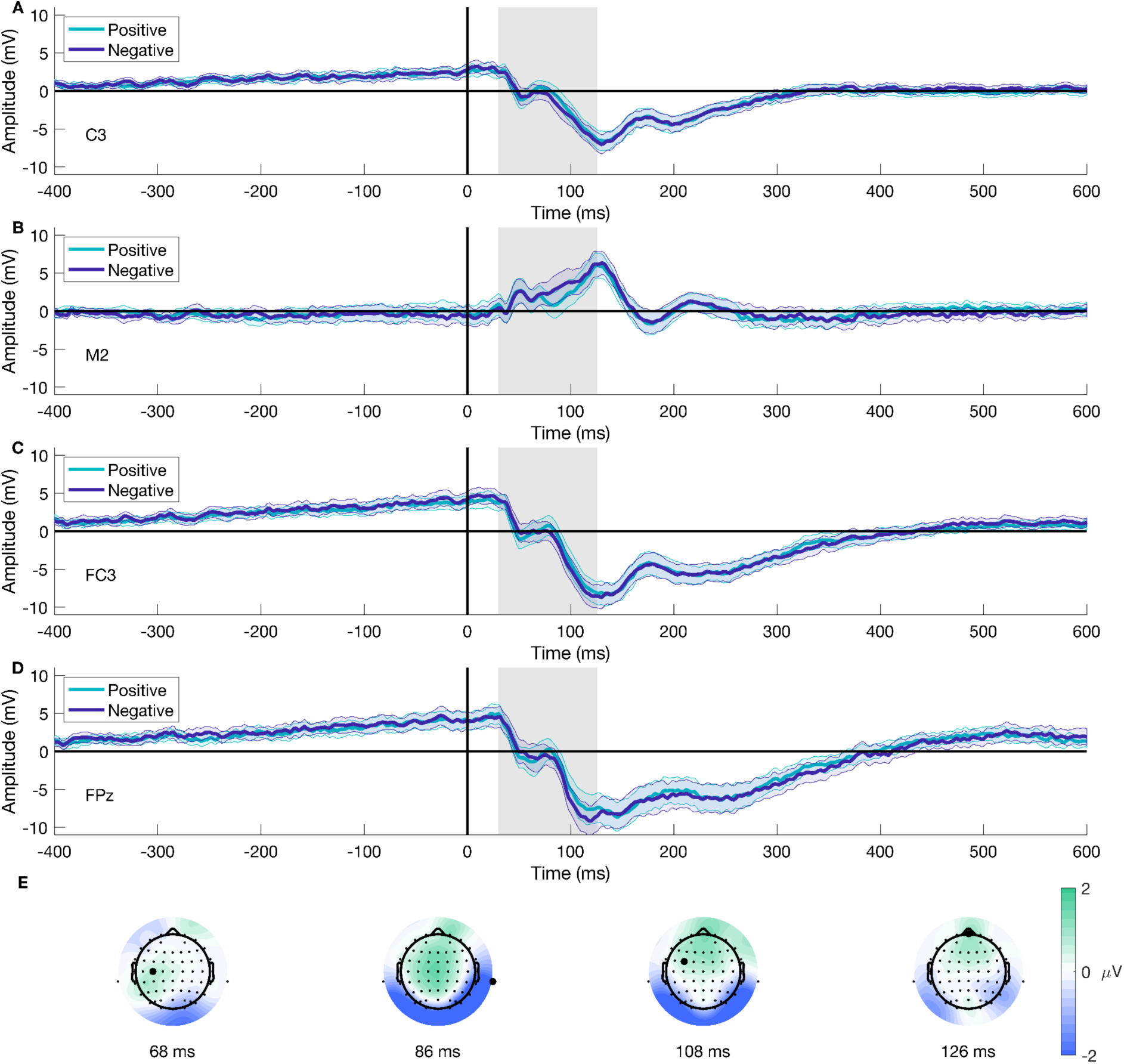
Stimulus-onset ERP for Valence and topographical distribution of the cluster over time. Panels A to D show the mean ERP for different electrodes contributing to the cluster for both positive and negative trials. Panel E displays the average difference between conditions over time at four distinct time points. The gray area represents the temporal extension of the cluster. Highlighted electrodes on E show the position of the ERP channels. Panel B and the second topographical map on Panel E are also shown in Figures 3 and 4, respectively. We have shortened the temporal window to -400 to 600 ms for visualization purposes.

Similarly, the mass univariate analysis showed differences in the interaction of factors on the picture onset ERPs. We found a significant cluster suitable with an effect spanning from 574 to 618 ms post-picture onset. The cluster encompassed a total of 15^5^ electrodes (Figure 6). We found the cluster peak at channel P6, at 582 ms after picture onset (Figure 6, third topoplot). Our results showed a late main effect due to differences between conditions in limited time and space windows.

**Figure 6.**
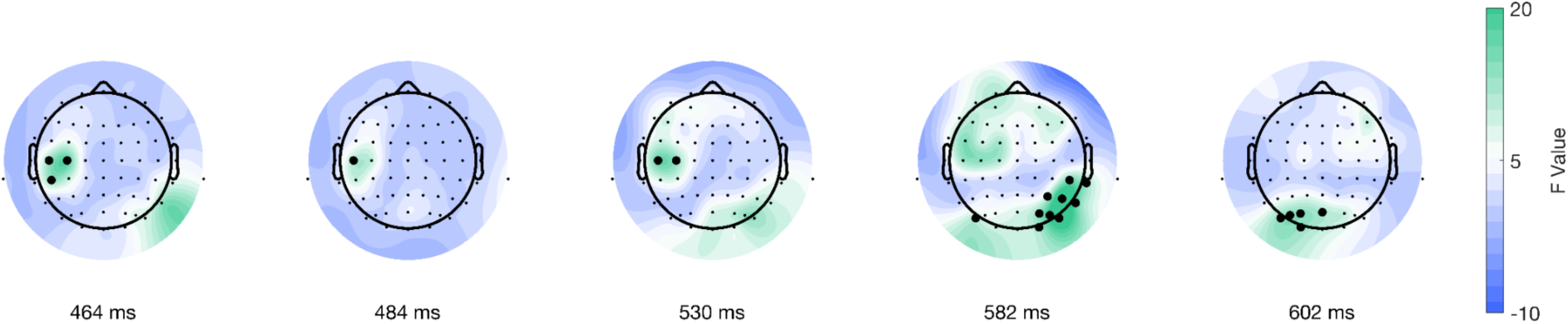
Topographical distribution of F values for the stimulus-locked interaction of factors as a function of time. The different topographical plots show the distribution of F values from the TFCE analysis over time. Highlighted electrodes represent those that are part of the significant cluster at each time point.

### Movement-Locked Cluster Characteristics

In the next sections, we will describe the cluster characteristics for the *Valence* factor and the interaction of factors for ERPs aligned to response-onset, given that there are no differences for the *Condition* factor.

Following the same procedure for the data epoched to stimulus-onset, we found differences in the *Valence* factor after performing a mass univariate analysis on the movement-onset epoched data. The analysis using TFCE (2×2 ANOVA, significance level < .05) showed a significant cluster for the *Valence* condition, corresponding with an effect spanning from -30 to 10 ms after movement onset (Figure 7A-D), which included 12^6^ out of the 64 electrodes. The peak of the cluster was on electrode P5, at 2 ms after movement onset, F(38) = 30.1562, p = 0.0056 (Figure 7C and the third difference plot in 7E), presenting a median difference between positive and negative trials of -1.3 µV (std = 1.483, [-5.342 2.284]) of this time-electrode combination. Furthermore, topographical dynamics are maintained through the cluster temporal window. This result shows significant early differences after movement onset regarding the emotional valence of the stimuli; the difference is driven by higher amplitudes observed during negative trials than positive ones.

**Figure 7.**
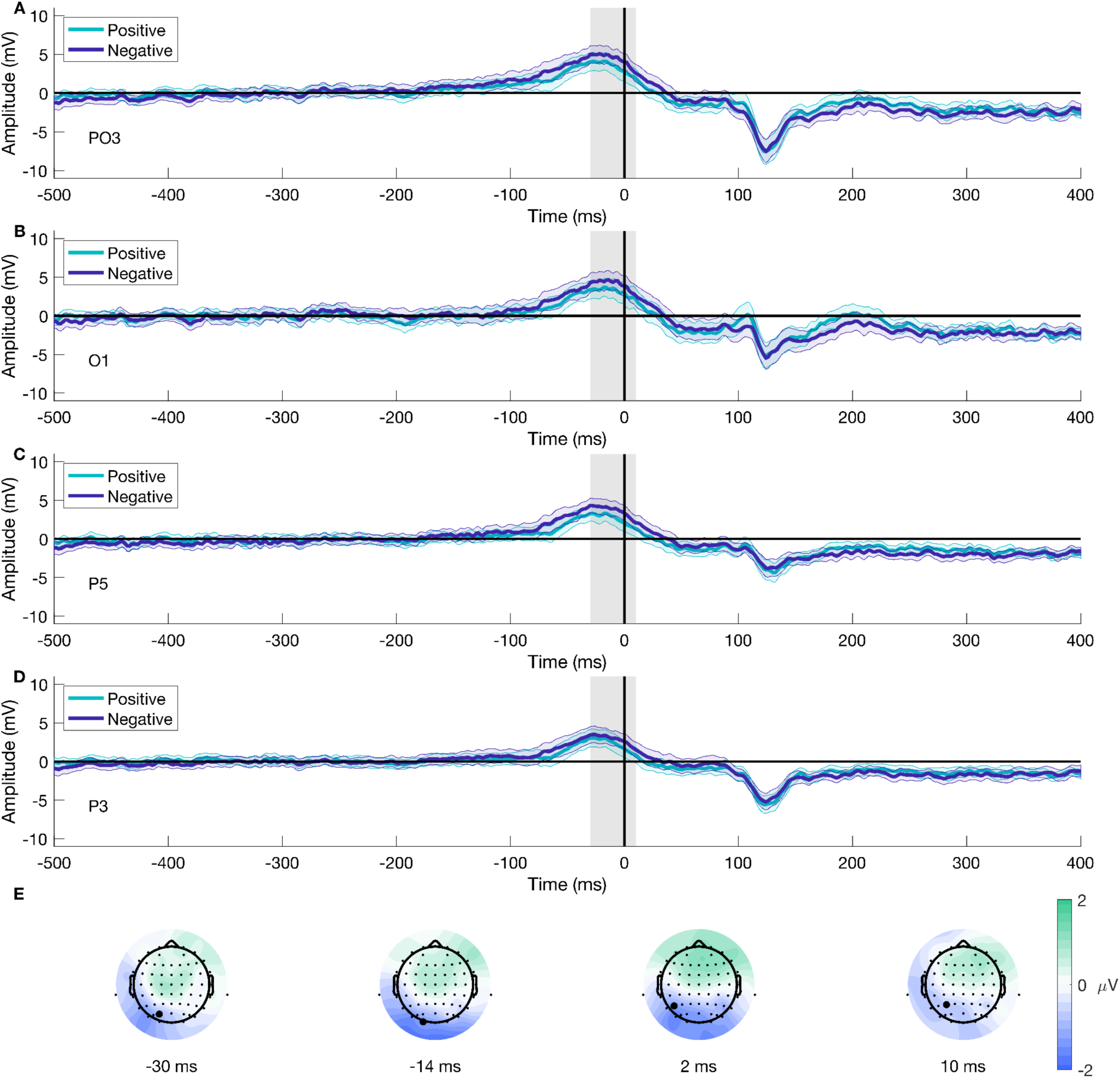
Movement-onset ERP for Valence and topographical distribution of the cluster over time. Panels A to D show the mean ERP for different electrodes contributing to the cluster for both positive and negative trials. Panel E displays the average difference between conditions over time at four distinct time points. The gray line represents the temporal peak of the cluster. Highlighted electrodes on E show the position of the ERP channels. Panel C and the third topographical map on Panel E are also shown in Figures 3 and 4, respectively. We have shortened the temporal window to -500 to 400 ms for visualization purposes.

Furthermore, the mass univariate analysis showed differences in the interaction of factors on the movement-onset ERPs. We found a significant cluster suitable with an effect spanning from 28 to 312 ms post-movement onset encompassing all electrodes (Figure 8). We identified the cluster peak at channel POz at 108 ms after picture onset, F(38) = 86.6177, p < .0001 (Figure 8, third topoplot). The topographical dynamics are maintained through the cluster temporal window. Overall, our results showed a late main effect due to differences between conditions in a relatively limited time window but a large spatial distribution encompassing all electrodes.

**Figure 8.**
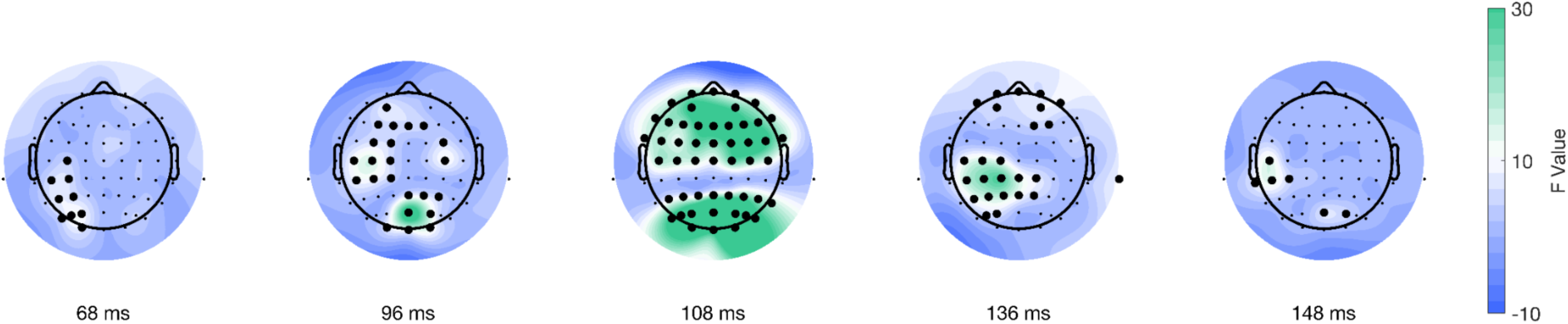
Topographical distribution of F values for the movement-locked interaction of factors as a function of time. The different topographical plots show the distribution of F values from the TFCE analysis over time. Highlighted electrodes represent those that are part of the significant cluster at each time point.

#### EEG Analysis: Frontal Alpha Asymmetry (FAA)

We conducted non-parametric tests (Wilcoxon Signed-Rank test) to assess differences in FAA after stimulus-onset. First, we computed FAA on single electrodes (F4 and F3) and assessed the difference between pleasant and unpleasant trials, which resulted in no significant differences (p = .961). Then, we computed FAA averaging frontal electrodes surrounding F4 (AF4, F2, F6, F4, FC2, FC4, FC6) and F3 (AF3, F5, F1, F3, FC3, FC5, FC1) on the 10-20 positioning system, to get a cluster approximation for frontal-right and frontal-left electrodes as described in the literature (Lacey & Gable, 2021; Vincent et al., 2021). As before, we assessed differences between pleasant and unpleasant trials, which showed no significant differences (p = .604). The results from the FAA analyses showed no significant differences in alpha asymmetry for pleasant and unpleasant trials.

## Discussion

The present study aimed to explore neural dynamics within the AAB. To this end, we replicated the classic AAT setup implemented by Solzbacher and colleagues (2022) and collected EEG data. Then, we analyzed the behavioral data and differences in the time-series domain and FAA of the neurophysiological data. We found significant differences in behavioral and neurophysiological data, which we will demonstrate in the following sections to partially align with and extend upon previous findings in the field.

Our behavioral results show a main effect on the *Valence* and *Condition* factors, replicating findings described in the literature and supporting the existence of the bias as assessed by the AATs. We found that reaction times were shorter when participants interacted with a pleasurable rather than an unpleasurable valenced picture. Similarly, we found that reaction times were shorter when participants performed a movement congruent with the natural predispositions (e.g., pushing the unpleasurable pictures and pulling the pleasurable ones) than reaction times for the incongruent conditions. However, we did not find significant differences in the interaction of factors. These results align with the findings described by Solzbacher and colleagues (2022) as well as in other studies (Czeszumski et al., 2021; Lang et al., 1990; Phaf et al., 2014) and support the assumptions about approach-avoidance behaviors. This reflects the evolutionary-rooted tendency to approach faster stimuli perceived as positive or beneficial rather than harmful or adverse ones and to be faster when behaving congruently than incongruently regarding natural predisposition.

Furthermore, the time-series analyses of movement and picture onset showed significant effects on the interaction of *Condition and Valence* and the *Valence* factor. Regarding the interactional effect, this result best describes the tendency related to the AAB (i.e., being faster to approach the positive and avoid the negative) and reflects the experimental design commonly implemented to study the bias. Hence, it proves the adequacy of the implemented behavioral task and analytical strategy for studying the neural dynamics of the AAB. For stimulus-locked trials, our results show a significant difference between conditions with a cluster from 574 to 618 ms (peak at 582 ms), with a parietal-posterior distribution. Previous studies on the AAB have reported late differences in the interaction of factors that align in time and topography with the Late Positive Potential (LPP; Bamford et al., 2015; Sege et al., 2024). However, our results show a negative potential, which challenges common interpretations about the temporality and spatial location of the difference and the corresponding underlying cognitive processes. The negativity, as well as the spatial and temporal resolution of the difference, point out the direction of the readiness potential (Colebatch, 2007; Shibasaki & Hallett, 2006), highlighting the period before movement onset where the decision regarding the movement to perform based on the emotional valence and the given instructions is made (Alexander et al., 2014; Lui et al., 2021; Schurger et al., 2021). Furthermore, response-locked trials revealed a significant cluster from 28 to 312 ms, with a maximum peak at 108 ms, located in the parietal-occipital region. We would have expected earlier negative differences, mostly previous movement onset, as a marker of the readiness potential for preparation to action. Nevertheless, to our knowledge, this study is one of the first attempts to systematically explore movement onset ERPs in the context of the AAB. Our cluster does not match the temporal characteristics of the readiness potential. We believe the difference we see can be explained by the task’s properties, such as the stimuli’s zooming effect, which aligns with our results’ temporal and spatial characteristics. In summary, we found significant differences in ERPs aligned to picture and movement onset regarding the interactional factor. Our results for stimulus-locked trials most likely correspond to differences regarding movement preparation and decision-making rather than aligning with previous findings. Additionally, they highlight the need for further analyses of response-locked ERPs.

Investigating the influence of valence revealed significant differences in ERPs for negative compared to positive stimuli in trials aligned to picture onset, temporally and spatially corresponding to the P100 component. These findings are consistent with previous research reporting differences in the P100 amplitude associated with emotional processing (Bublatzky & Schupp, 2012; Gerdes et al., 2013; Pourtois et al., 2004). These differences are believed to reflect the early stages of emotional processing (Ding et al., 2017; Pourtois et al., 2004), potentially priming the visual signal to timely responses (Bublatzky & Schupp, 2012; Smith et al., 2003). Response-locked trials revealed a significant cluster centered around or shortly after movement onset. To our knowledge, our study is the first to investigate valence-related differences for movement-related ERPs in the AAB paradigm. The underlying neuronal processes contributing to the observed cluster are not identifiable based on our results alone. The cluster’s peak was contralateral, suggesting that early stages of movement execution likely contributed to the observed differences (Kornhuber & Deecke, 2016). Moreover, the cluster extended to before movement onset, consistent with the readiness potential (Colebatch, 2007; Shibasaki & Hallett, 2006). However, the positive rather than negative potential challenges its interpretation and the attribution to readiness potential differences. Further research is needed to disentangle the contributions of valence to movement-related ERP differences. Overall, we observed differences in response-locked and stimulus-locked trials for the valence factor. These results support the existence of early differences in the P100 component and are the first to investigate variations associated with movement execution.

Unexpectedly, our study did not find a significant effect of the *Condition* factor. Considering the relevance of deliberation and reflective reasoning processes based on cognitive control functions (Loijen et al., 2020), we would have expected to find differences in brain activity when comparing congruent and incongruent conditions. This is mainly due to the high cognitive load of counter-intuitive decisions (e.g., approaching an unpleasant picture or avoiding a pleasant one Jackson et al., 2014; Zhou et al., 2017). Furthermore, given the role of the PFC as part of both the underlying mechanisms for approach and avoidance behaviors, as well as for decision-making processes (Ironside et al., 2020; Kaldewaij et al., 2017; Lender et al., 2023; Radke et al., 2015; Volman et al., 2011; Wiers et al., 2014; Zorowitz et al., 2019), we would have expected to see differences located in frontal regions either after picture onset or before movement onset, which will correspond to the deliberation or decision-making period. Even when some studies have reported differences in the *Condition* factor when analyzing ERPs in different contexts of the AAB (Cheval et al., 2018; Ernst et al., 2013; Marrero et al., 2023), it might be that due to the characteristics of the primary structure involved i.e., the amygdala (Andrewes & Jenkins, 2019; Loijen et al., 2020; Ochsner & Gross, 2005; Sergerie et al., 2008; Wager et al., 2008), that a cortical analysis such as ERP or EEG, in general, is not a suited approximation and other techniques that can provide better insights for subcortical analyses should be considered (e.g., fMRI or functional Near Infrared Spectroscopy; fNIRS). Similarly, two methodological aspects of the design might impact our result. First, the open temporal window for reaction time can include different cognitive processes, especially when considering the differences in reaction times (Table 1 and Behavioral Results Section). Secondly, the reduced number of trials (N = 20 per condition) might not be enough to elicit an ERP and account for the statistical power of the differences. While the current study did not find the expected effect of the *Condition* factor, employing different methodological approaches in future studies might better describe the brain dynamics related to the AAB.

In contrast to the time-series analysis, our results showed no significant differences between FAA values for pleasant and unpleasant trials for stimulus-locked trials. Higher left hemispheric activity tends to be associated with positive and pleasurable stimuli, while higher right activity has been associated with negative valenced stimuli (Briesemeister et al., 2013; Sabu et al., 2022). Given the characteristics of FAA, it has been a preferred measure to explore activity related to affective stimuli and motivation (Briesemeister et al., 2013; Sabu et al., 2022); nevertheless, results are heterogeneous. The review conducted by Sabu and colleagues (2022) shows that out of 18 studies using emotionally valenced pictures to assess differences in FAA, only two found an effect of the presented stimuli (Huster et al., 2009; Schöne et al., 2016). However, studies implementing videos (Papousek et al., 2014; Zhao et al., 2018), real cues (Knott et al., 2008; Olszewska-Guizzo et al., 2020), and games (Miller & Tomarken, 2001; Rodrigues et al., 2018) were more prone to find FAA differences. These results highlight a relevant methodological point regarding the suitability of static images for generating emotional engagement in the subjects and eliciting differences in FAA activity, especially when considering the positive results found by implementing relatable real-world cues or more immersive experiences by using videos, games, and even 3D stimuli and virtual environments (Grasso-Cladera et al., 2024; Sabu et al., 2022; Schöne, 2022). The heterogeneity in FAA results regarding emotionally valenced stimuli, and the high prevalence of negative results when using static stimuli or 2D pictures posits the need for new methodological approaches for studying the AAB under more emotionally engaging and naturalistic scenarios.

## Conclusion

This study systematically explored neural dynamics within the Attentional Bias (AAB) by implementing a classic AAT setup and collecting both behavioral and neurophysiological data. Our exploratory analysis revealed significant differences between the behavioral data and ERP findings, though no significant differences were found for the FAA. Our behavioral results align with existing literature on the AAB, which adds to the extensive body of evidence supporting the evolutive mechanisms of the bias. In contrast, the ERP results offer partial support for previous studies, though the literature on the neural dynamics of the AAB remains varied and heterogeneous. Importantly, this study is the first to systematically assess differences across time and topographies for both stimulus- and response-locked ERPs, marking a significant contribution to the field. We aim for this work to serve as a foundation for further systematic investigations into the neural dynamics of the AAB, with the potential to deepen our understanding of these processes. The absence of significant differences in FAA results is consistent with a substantial body of prior research, suggesting that methodological factors may play a key role in shaping these outcomes. Overall, our findings underscore the complexity of the neural mechanisms underlying the AAB and highlight the need for more nuanced methodologies in future studies. Despite the variability in the ERP results, our study offers important insights into the temporal and topographical patterns of neural activity associated with the AAB. Further research should build upon these findings, refining experimental paradigms and employing more sophisticated analytical techniques to clarify the neural dynamics of attentional biases.

## Data Availability Statement

The raw data is publicly available on the Open Science Platform: https://osf.io/4fmw9/

## Ethics Approval Statement

The study was approved by the Ethics Committee of Osnabrück University. Participants provided their written informed consent to participate in this study.

## Authors Contribution

JS and PK conceptualized and designed the experiment. JS carried out and supervised data collection. AGC preprocessed the data. AGC and DN conducted the analyses. AGC and DN wrote the first draft of the manuscript.

## Acknowledgments

The authors would like to thank Julian Webb for programming the experimental task and Johanna Kopetsch for her work on the preliminary EEG preprocessing and ERP analysis.

## Code availability

The code use for the analysis of the present article is available at: https://github.com/AitanaGrasso-Cladera/Exploring-brain-dynamics-within-the-Approach-Avoidance-Bias

(Grasso-Cladera, A., & Nolte, D. Exploring-brain-dynamics-within-the-Approach-Avoidance-Bias (Version 1.0) [Computer software]. https://github.com/AitanaGrasso-Cladera/Exploring-brain-dynamics-within-the-Approach-Avoidance-Bias)

---

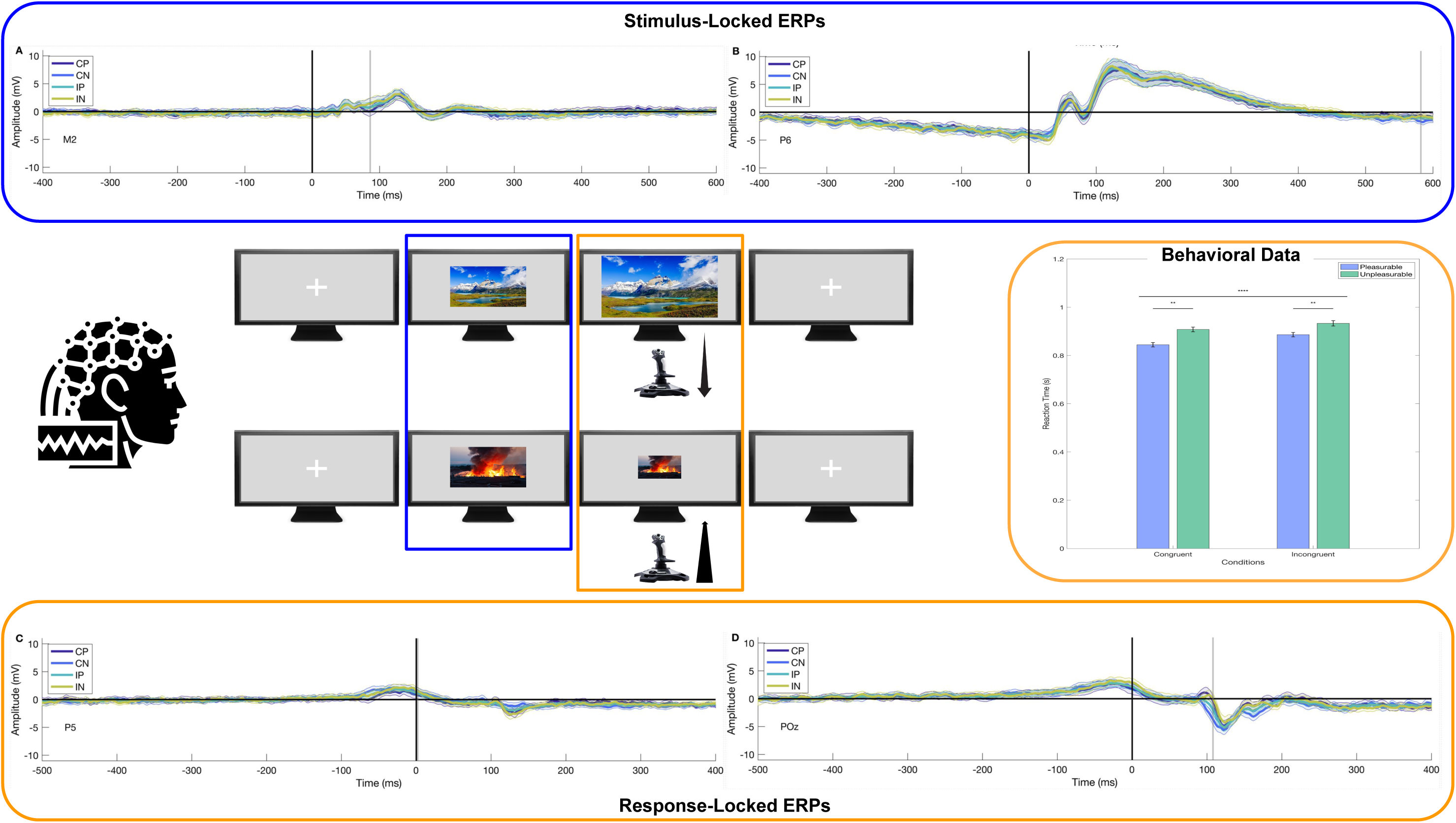

We systematically investigated stimulus (blue)- and response (orange)-locked event-related potentials and frontal alpha asymmetry for the Approach-Avoidance Bias (AAB). While our behavioral results confirmed the classic AAB effect, EEG analyses revealed early valence-related effects and late parietal-occipital differences but no significant differences in response congruency or frontal alpha asymmetry.

## Footnotes

1 Unpleasant pictures IDs: 2053, 2301, 2345.1, 2456, 2751, 3181, 3230, 3300, 6350, 6540, 6550, 6560, 6563, 6838, 9000, 9075, 9140, 9184, 9220, 9250, 9254, 9332, 9340, 9342, 9414, 9419, 9424, 9427, 9520, 9530, 9560, 9630, 9800, 9810, 9830, 9832, 9900, 9902, 9905, 9908, 9909, 9925, 9940, 9941.

2 Pleasant pictures IDs: 1340, 1410, 1500, 1540, 1600, 1610, 1620, 2040, 2057, 2058, 2075, 2150, 2158, 2160, 2170, 2208, 2260, 2274, 2299, 2300, 2306, 2314, 2332, 2340, 2341, 2345, 2347, 2360, 2387, 2388, 2391, 2392, 2398, 2540, 2598, 4599, 4614, 4628, 4640, 4641, 5200, 5202, 5210, 8497.

3 In order to maintain the experimental value of the IAPS collection, the images displayed in the figure are not part of it, but randomly selected pictures for the graphical representation of the experimental paradigm. Panel A: Sriwongthai, (n.d.); Panel B: Mulder, (n.d.).

4 Channels included in the cluster: AF3, AF4, AF7, AF8, C1, C2, C3, C4, C5, CP1, CP2, CP3, CP5, CP6, CPz, Cz, F1, F2, F3, F4, F5, F6, F8, FC1, FC2, FC3, FC4, FC6, FCz, FP1, FP2, FPz, Fz, M1, M2, O1, O2, Oz, P1, P2, P3, P4, P5, P6, P7, P8, PO3, PO4, PO5, PO6, PO7, PO8, POz, Pz, T8, TP7, TP8

5 Channels included in the cluster: CP6, O1, O2, Oz, P4, P6, P8, PO3, PO4, PO5, PO6, PO7, PO8, POz, TP8

6 Channels included in the cluster: CP5, O1, O2, Oz, P1, P3, P5, P7, PO3, PO5, PO7, TP7

## References

Alexander, P., Schlegel, A. A., Sinnott-Armstrong, W., Roskies, A., Tse, P., & Wheatley, T. (2014). Dissecting the readiness potential. Surrounding Free Will: Philosophy, Psychology, Neuroscience, 203–230.

Andrewes, D. G., & Jenkins, L. M. (2019). The Role of the Amygdala and the Ventromedial Prefrontal Cortex in Emotional Regulation: Implications for Post-traumatic Stress Disorder. Neuropsychology Review, 29(2), 220–243.

Aupperle, R. L., Melrose, A. J., Francisco, A., Paulus, M. P., & Stein, M. B. (2015). Neural substrates of approach-avoidance conflict decision-making. Human Brain Mapping, 36(2), 449–462.

Bamford, S., Broyd, S. J., Benikos, N., Ward, R., Wiersema, J. R., & Sonuga-Barke, E. (2015). The late positive potential: a neural marker of the regulation of emotion-based approach-avoidance actions? Biological Psychology, 105, 115–123.

Bigdely-Shamlo, N., Mullen, T., Kothe, C., Su, K.-M., & Robbins, K. A. (2015). The PREP pipeline: standardized preprocessing for large-scale EEG analysis. Frontiers in Neuroinformatics, 9, 16.

Bradley, M., Codispoti, M., Cuthbert, B., & Lang, P. (2001). Emotion and motivation I: defensive and appetitive reactions in picture processing. Emotion (Washington, D.C.), 1(3), 276–298.

Branco, D., Gonçalves, Ó. F., & Badia, S. B. (2023). A Systematic Review of International Affective Picture System (IAPS) around the World. Sensors . https://www.mdpi.com/1424-8220/23/8/3866

Briesemeister, B. B., Tamm, S., Heine, A., & Jacobs, A. (2013). Approach the good, withdraw from the bad—A review on frontal alpha asymmetry measures in applied psychological research. Psychology, 4, 261–267.

Bublatzky, F., & Schupp, H. T. (2012). Pictures cueing threat: brain dynamics in viewing explicitly instructed danger cues. Social Cognitive and Affective Neuroscience, 7(6), 611–622.

Chen, M., & Bargh, J. A. (1999). Consequences of Automatic Evaluation: Immediate Behavioral Predispositions to Approach or Avoid the Stimulus. Personality & Social Psychology Bulletin, 25(2), 215–224.

Cheval, B., Tipura, E., Burra, N., Frossard, J., Chanal, J., Orsholits, D., Radel, R., & Boisgontier, M. P. (2018). Avoiding sedentary behaviors requires more cortical resources than avoiding physical activity: An EEG study. Neuropsychologia, 119, 68–80.

Cohen, J. (1988). Statistical power analysis for the behavioral sciences (2nd ed.). Lawrence Erlbaum Associates. https://www.utstat.toronto.edu/~brunner/oldclass/378f16/readings/CohenPower.pdf

Colebatch, J. G. (2007). Bereitschaftspotential and movement-related potentials: origin, significance, and application in disorders of human movement. Movement Disorders: Official Journal of the Movement Disorder Society, 22(5), 601–610.

Czeszumski, A., Albers, F., Walter, S., & König, P. (2021). Let Me Make You Happy, and I’ll Tell You How You Look Around: Using an Approach-Avoidance Task as an Embodied Emotion Prime in a Free-Viewing Task. Frontiers in Psychology, 12, 604393.

de Cheveigné, A. (2020). ZapLine: A simple and effective method to remove power line artifacts. NeuroImage, 207, 116356.

Degner, J., Steep, L., Schmidt, S., & Steinicke, F. (2021). Assessing Automatic Approach-Avoidance Behavior in an Immersive Virtual Environment. Frontiers in Virtual Reality, 2. 10.3389/frvir.2021.761142

Ding, R., Li, P., Wang, W., & Luo, W. (2017). Emotion processing by ERP combined with development and plasticity. Neural Plasticity, 2017, 5282670.

Durlak, J. (2009). How to select, calculate, and interpret effect sizes. Journal of Pediatric Psychology, 34(9), 917–928.

Ehinger, B. V., & Dimigen, O. (2018). Unfold: an integrated toolbox for overlap correction, non-linear modeling, and regression-based EEG analysis. bioRxiv, 7. 10.7717/peerj.7838

Eiler, T. J., Grünewald, A., Machulska, A., Klucken, T., Jahn, K., Niehaves, B., Gethmann, C. F., & Brück, R. (2019). A Preliminary Evaluation of Transferring the Approach Avoidance Task into Virtual Reality. Information Technology in Biomedicine, 151–163.

Elliot, A. J. (2006). The Hierarchical Model of Approach-Avoidance Motivation. Motivation and Emotion, 30(2), 111–116.

Ellis, A. J., Salgari, G., Miklowitz, D. J., & Loo, S. K. (2019). The role of avoidance motivation in the relationship between reward sensitivity and depression symptoms in adolescents: An ERP study. Psychiatry Research, 279, 345–349.

Ernst, L. H., Ehlis, A.-C., Dresler, T., Tupak, S. V., Weidner, A., & Fallgatter, A. J. (2013). N1 and N2 ERPs reflect the regulation of automatic approach tendencies to positive stimuli. Neuroscience Research, 75(3), 239–249.

Fechtner, J. (2013). The Role of Cognitive Control and Approach-Avoidance Motivation in the Relationship between Stress and Aggression-A Psychophysiological Investigation. https://ubt.opus.hbz-nrw.de/opus45-ubtr/frontdoor/index/index/docId/561

Fox, N. A., Rubin, K. H., Calkins, S. D., Marshall, T. R., Coplan, R. J., Porges, S. W., Long, J. M., & Stewart, S. (1995). Frontal activation asymmetry and social competence at four years of age. Child Development, 66(6), 1770–1784.

Fridland, E., & Wiers, C. E. (2018). Addiction and embodiment. Phenomenology and the Cognitive Sciences, 17(1), 15–42.

Gerdes, A. B. M., Wieser, M. J., Bublatzky, F., Kusay, A., Plichta, M. M., & Alpers, G. W. (2013). Emotional sounds modulate early neural processing of emotional pictures. Frontiers in Psychology, 4, 741.

Goulet-Pelletier, J. C., & Cousineau, D. (2018). A review of effect sizes and their confidence intervals, Part I: The Cohen’sd family. The Quantitative Methods for Psychology, 14(4), 242–265.

Grasso-Cladera, A., Madrid-Carvajal, J., Walter, S., & König, P. (2024). Approach-Avoidance Bias in Virtual and Real-World Simulations: Insights from a Systematic Review of Experimental Setups. 2024.12. 12.628144.

Huster, R. J., Stevens, S., Gerlach, A. L., & Rist, F. (2009). A spectralanalytic approach to emotional responses evoked through picture presentation. International Journal of Psychophysiology: Official Journal of the International Organization of Psychophysiology, 72(2), 212–216.

Ironside, M., Amemori, K.-I., McGrath, C. L., Pedersen, M. L., Kang, M. S., Amemori, S., Frank, M. J., Graybiel, A. M., & Pizzagalli, D. A. (2020). Approach-Avoidance Conflict in Major Depressive Disorder: Congruent Neural Findings in Humans and Nonhuman Primates. Biological Psychiatry, 87(5), 399–408.

Jackson, S. A., Kleitman, S., & Aidman, E. (2014). Low cognitive load and reduced arousal impede practice effects on executive functioning, metacognitive confidence and decision making. PloS One, 9(12), e115689.

Jain, A., Bansal, R., Kumar, A., & Singh, K. D. (2015). A comparative study of visual and auditory reaction times on the basis of gender and physical activity levels of medical first year students. International Journal of Applied & Basic Medical Research, 5(2), 124–127.

Kaldewaij, R., Koch, S. B. J., Volman, I., Toni, I., & Roelofs, K. (2017). On the Control of Social Approach–Avoidance Behavior: Neural and Endocrine Mechanisms. In M. Wöhr & S. Krach (Eds.), Social Behavior from Rodents to Humans: Neural Foundations and Clinical Implications (pp. 275–293). Springer International Publishing.

Kaspar, K., Gameiro, R. R., & König, P. (2015). Feeling good, searching the bad: Positive priming increases attention and memory for negative stimuli on webpages. Computers in Human Behavior, 53, 332–343.

Kenrick, D. T., & Shiota, M. N. (2008). Approach and avoidance motivation(s): An evolutionary perspective. In A. J. Elliot (Ed.), Handbook of approach and avoidance motivation*, (pp* (Vol. 664, pp. 273–288). Psychology Press, xvii.

Kleiner, M., Brainard, D., & Pelli, D. (2007). What’s new in Psychtoolbox-3? https://pure.mpg.de/rest/items/item_1790332/component/file_3136265/content

Klug, M., & Kloosterman, N. A. (2022). Zapline-plus: A Zapline extension for automatic and adaptive removal of frequency-specific noise artifacts in M/EEG. Human Brain Mapping. 10.1002/hbm.25832

Knott, V. J., Naccache, L., Cyr, E., Fisher, D. J., McIntosh, J. F., Millar, A. M., & Villeneuve, C. M. (2008). Craving-induced EEG reactivity in smokers: effects of mood induction, nicotine dependence and gender. Neuropsychobiology, 58(3-4), 187–199.

Kornhuber, H. H., & Deecke, L. (2016). Brain potential changes in voluntary and passive movements in humans: readiness potential and reafferent potentials. Pflugers Archiv: European Journal of Physiology, 468(7), 1115–1124.

Kothe, C. (2014). Lab streaming layer (LSL). https://github.Com/sccn/labstreaminglayer. Accessed on October, 26, 2015.

Krieglmeyer, R., De Houwer, J., & Deutsch, R. (2013). On the nature of automatically triggered approach–avoidance behavior. Emotion Review: Journal of the International Society for Research on Emotion, 5(3), 280–284.

Lacey, M. F., & Gable, P. A. (2021). Frontal Asymmetry in an approach-avoidance conflict paradigm. Psychophysiology, 58(5), e13780.

Lang, P. J. (1995). The emotion probe. Studies of motivation and attention. The American Psychologist, 50(5), 372–385.

Lang, P. J., Bradley, M. M., & Cuthbert, B. N. (1990). Emotion, attention, and the startle reflex. Psychological Review, 97(3), 377–395.

Lang, P. J., Bradley, M. M., & Cuthbert, B. N. (1997). International affective picture system (IAPS): Technical manual and affective ratings. NIMH Center for the Study of. https://www2.unifesp.br/dpsicobio/adap/instructions.pdf

Lang, P. J., Bradley, M. M., Fitzsimmons, J. R., Cuthbert, B. N., Scott, J. D., Moulder, B., & Nangia, V. (1998). Emotional arousal and activation of the visual cortex: an fMRI analysis. Psychophysiology, 35(2), 199–210.

Lang, P. J., Greenwald, M. K., Bradley, M. M., & Hamm, A. O. (1993). Looking at pictures: affective, facial, visceral, and behavioral reactions. Psychophysiology, 30(3), 261–273.

Lender, A., Wirtz, J., Kronbichler, M., Kahveci, S., Kühn, S., & Blechert, J. (2023). Differential Orbitofrontal Cortex Responses to Chocolate Images While Performing an Approach– Avoidance Task in the MRI Environment. Nutrients, 15(1), 244.

Loijen, A., Vrijsen, J. N., Egger, J. I. M., Becker, E. S., & Rinck, M. (2020). Biased approach-avoidance tendencies in psychopathology: A systematic review of their assessment and modification. Clinical Psychology Review, 77, 101825.

Lui, K. K., Nunez, M. D., Cassidy, J. M., Vandekerckhove, J., Cramer, S. C., & Srinivasan, R. (2021). Timing of readiness potentials reflect a decision-making process in the human brain. Computational Brain & Behavior, 4(3), 264–283.

Marrero, H., Yagual, S. N., Lemus, A., García-Marco, E., Díaz, J. M., Gámez, E., Urrutia, M., & Beltrán, D. (2023). Social approach and avoidance in language: N400-like ERP negativity indexes congruency and theta rhythms the conflict. Cerebral Cortex (New York, N.Y.: 1991), 33(4), 1300–1309.

McManis, M. H., Bradley, M. M., Berg, W. K., Cuthbert, B. N., & Lang, P. J. (2001). Emotional reactions in children: verbal, physiological, and behavioral responses to affective pictures. Psychophysiology, 38(2), 222–231.

McNaughton, N., DeYoung, C. G., & Corr, P. J. (2016). Chapter 2 - Approach/Avoidance. In J. R. Absher & J. Cloutier (Eds.), Neuroimaging Personality, Social Cognition, and Character (pp. 25–49). Academic Press.

Mensen, A., & Khatami, R. (2013). Advanced EEG analysis using threshold-free cluster-enhancement and non-parametric statistics. NeuroImage, 67, 111–118.

Miller, A., & Tomarken, A. J. (2001). Task-dependent changes in frontal brain asymmetry: Effects of incentive cues, outcome expectancies, and motor responses. Psychophysiology, 38(3), 500–511.

Morís Fernández, L., & Vadillo, M. A. (2020). Flexibility in reaction time analysis: many roads to a false positive? Royal Society Open Science, 7(2), 190831.

Mulder, Vladimir. (n.d.). Large blaze at burning industrial building [Stock image]. Shutterstock. https://www.shutterstock.com/image-photo/large-blaze-burning-industrial-building-2489081119

Ochsner, K. N., & Gross, J. J. (2005). The cognitive control of emotion. Trends in Cognitive Sciences, 9(5), 242–249.

Olszewska-Guizzo, A., Sia, A., Fogel, A., & Ho, R. (2020). Can exposure to certain urban green spaces trigger frontal alpha asymmetry in the brain?-preliminary findings from a passive task EEG study. International Journal of Environmental Research and Public Health, 17(2), 394.

Otaki, M., & Shibata, K. (2019). The effect of different visual stimuli on reaction times: a performance comparison of young and middle-aged people. Journal of Physical Therapy Science, 31, 250–254.

Palmer, J. A., Kreutz-Delgado, K., & Makeig, S. (2012). AMICA: An adaptive mixture of independent component analyzers with shared components. Swartz Center for Computatonal Neursoscience, University of California San Diego, Tech. Rep. https://www.academia.edu/download/46689710/AMICA_An_Adaptive_Mixture_of_Independent20160621-888-19z4hv3.pdf

Papousek, I., Weiss, E. M., Schulter, G., Fink, A., Reiser, E. M., & Lackner, H. K. (2014). Prefrontal EEG alpha asymmetry changes while observing disaster happening to other people: cardiac correlates and prediction of emotional impact. Biological Psychology, 103, 184–194.

Park, H.-B., & Hyun, J.-S. (2014). The ex-Gaussian analysis of reaction time distributions for cognitive experiments. Science of Emotion and Sensibility, 17, 63–76.

Parsons, S., Kruijt, A.-W., & Fox, E. (2016). A Cognitive Model of Psychological Resilience. Journal of Experimental Psychopathology, 7(3), 296–310.

Phaf, R. H., Mohr, S. E., Rotteveel, M., & Wicherts, J. M. (2014). Approach, avoidance, and affect: a meta-analysis of approach-avoidance tendencies in manual reaction time tasks. Frontiers in Psychology, 5, 378.

Pourtois, G., Grandjean, D., Sander, D., & Vuilleumier, P. (2004). Electrophysiological correlates of rapid spatial orienting towards fearful faces. Cerebral Cortex (New York, N.Y.: 1991), 14(6), 619–633.

Radke, S., Volman, I., Mehta, P., van Son, V., Enter, D., Sanfey, A., Toni, I., de Bruijn, E. R. A., & Roelofs, K. (2015). Testosterone biases the amygdala toward social threat approach. Science Advances, 1(5), e1400074.

Ramnani, N., & Owen, A. M. (2004). Anterior prefrontal cortex: insights into function from anatomy and neuroimaging. Nature Reviews. Neuroscience, 5(3), 184–194.

Renton, A. I., Painter, D. R., & Mattingley, J. B. (2019). Differential Deployment of Visual Attention During Interactive Approach and Avoidance Behavior. Cerebral Cortex , 29(6), 2366–2383.

Rodrigues, J., Müller, M., Mühlberger, A., & Hewig, J. (2018). Mind the movement: Frontal asymmetry stands for behavioral motivation, bilateral frontal activation for behavior. Psychophysiology, 55(1). 10.1111/psyp.12908

Roelofs, K., van Peer, J., Berretty, E., Jong, P. de, Spinhoven, P., & Elzinga, B. M. (2009). Hypothalamus–Pituitary–Adrenal Axis Hyperresponsiveness Is Associated with Increased Social Avoidance Behavior in Social Phobia. Biological Psychiatry, 65(4), 336–343.

Rousselet, G. A., & Pernet, C. R. (2011). Quantifying the time course of visual object processing using ERPs: It’s time to up the game. Frontiers in Psychology, 2, 107.

Rousselet, G. A., & Wilcox, R. R. (2018). Reaction times and other skewed distributions: problems with the mean and the median. bioRxiv. 10.1101/383935

Sabu, P., Stuldreher, I. V., Kaneko, D., & Brouwer, A.-M. (2022). A review on the role of affective stimuli in event-related frontal alpha asymmetry. Frontiers in Computer Science, 4. 10.3389/fcomp.2022.869123

Schmidt, V., & Nolte, D. (2024). EEG_preprocessing_scripts [MATLAB]. https://github.com/debnolte/EEG_preprocessing_scripts

Schöne, B. (2022). Commentary: A review on the role of affective stimuli in event-related frontal alpha asymmetry. Frontiers in Computer Science, 4. 10.3389/ fcomp.2022.994071

Schöne, B., Schomberg, J., Gruber, T., & Quirin, M. (2016). Event-related frontal alpha asymmetries: electrophysiological correlates of approach motivation. Experimental Brain Research, 234(2), 559–567.

Schurger, A., Hu, P. ‘ben’, Pak, J., & Roskies, A. L. (2021). What is the readiness potential? Trends in Cognitive Sciences, 25(7), 558–570.

Sege, C. T., Lopez, J. W., Hellman, N. M., & McTeague, L. M. (2024). Assessing motivational biases in brain and behavior: Event-related potential and response time concomitants of the approach-avoidance task. Psychophysiology, 61(12), e14700.

Sergerie, K., Chochol, C., & Armony, J. L. (2008). The role of the amygdala in emotional processing: a quantitative meta-analysis of functional neuroimaging studies. Neuroscience and Biobehavioral Reviews, 32(4), 811–830.

Shibasaki, H., & Hallett, M. (2006). What is the bereitschaftspotential? Clinical Neurophysiology: Official Journal of the International Federation of Clinical Neurophysiology, 117(11), 2341–2356.

Smith, N. K., Cacioppo, J. T., Larsen, J. T., & Chartrand, T. L. (2003). May I have your attention, please: electrocortical responses to positive and negative stimuli. Neuropsychologia, 41(2), 171–183.

Solarz, A. K. (1960). Latency of instrumental responses as a function of compatibility with the meaning of eliciting verbal signs. Journal of Experimental Psychology, 59, 239–245.

Solzbacher, J., Czeszumski, A., Walter, S., & König, P. (2022). Evidence for the embodiment of the automatic approach bias. Frontiers in Psychology, 13, 797122.

Sriwongthai, Ichy. (n.d.). Beautiful Patagonia landscape of Andes mountain range, winding road and lake at Torres del Paine National Park, Chile. [Stock image]. Shutterstock. https://www.shutterstock.com/image-photo/beautiful-patagonia-landscape-andes-mountain-range-1172916133?consentChanged=true

Strack, F., & Deutsch, R. (2004a). Reflection and impulse as determinants of conscious and unconscious motivation. Social Motivation: Conscious and Unconscious Processes, 91–112.

Strack, F., & Deutsch, R. (2004b). Reflective and impulsive determinants of social behavior. Personality and Social Psychology Review: An Official Journal of the Society for Personality and Social Psychology, Inc, 8(3), 220–247.

Stropahl, M., & Debener, S. (2017). Auditory cross-modal reorganization in cochlear implant users indicates audio-visual integration. NeuroImage. Clinical, 16, 514–523.

Te Grotenhuis, M., Pelzer, B., Eisinga, R., Nieuwenhuis, R., Schmidt-Catran, A., & Konig, R. (2017). When size matters: advantages of weighted effect coding in observational studies. International Journal of Public Health, 62(1), 163–167.

Van Dessel, P., De Houwer, J., Gast, A., Smith, C. T., & De Schryver, M. (2016). Instructing implicit processes: When instructions to approach or avoid influence implicit but not explicit evaluation. Journal of Experimental Social Psychology, 63, 1–9.

Vecchio, A., & Pascalis, V. (2020). EEG resting asymmetries and frequency oscillations in approach/avoidance personality traits: A systematic review. Symmetry, 12, 1712.

Vincent, K. M., Xie, W., & Nelson, C. A. (2021). Using different methods for calculating frontal alpha asymmetry to study its development from infancy to 3 years of age in a large longitudinal sample. Developmental Psychobiology, 63(6), e22163.

Volman, I., Toni, I., Verhagen, L., & Roelofs, K. (2011). Endogenous Testosterone Modulates Prefrontal–Amygdala Connectivity during Social Emotional Behavior. Cerebral Cortex , 21(10), 2282–2290.

Wager, T. D., Davidson, M. L., Hughes, B. L., Lindquist, M. A., & Ochsner, K. N. (2008). Prefrontal-subcortical pathways mediating successful emotion regulation. Neuron, 59(6), 1037–1050.

Widmann, A., Schröger, E., & Maess, B. (2015). Digital filter design for electrophysiological data--a practical approach. Journal of Neuroscience Methods, 250, 34–46.

Wiers, C. E., Stelzel, C., Park, S. Q., Gawron, C. K., Ludwig, V. U., Gutwinski, S., Heinz, A., Lindenmeyer, J., Wiers, R. W., Walter, H., & Bermpohl, F. (2014). Neural correlates of alcohol-approach bias in alcohol addiction: the spirit is willing but the flesh is weak for spirits. Neuropsychopharmacology: Official Publication of the American College of Neuropsychopharmacology, 39(3), 688–697.

Winward, S. B., Siklos-Whillans, J., & Itier, R. J. (2022). Impact of face outline, parafoveal feature number and feature type on early face perception in a gaze-contingent paradigm: A mass-univariate re-analysis of ERP data. Neuroimage. Reports, 2(4), 100148.

Zech, H. G., Rotteveel, M., van Dijk, W. W., & van Dillen, L. F. (2020). A mobile approach-avoidance task. Behavior Research Methods, 52(5), 2085–2097.

Zhao, G., Zhang, Y., Ge, Y., Zheng, Y., Sun, X., & Zhang, K. (2018). Asymmetric hemisphere activation in tenderness: evidence from EEG signals. Scientific Reports, 8(1), 8029.

Zhou, J., Arshad, S. Z., Luo, S., & Chen, F. (2017). Effects of uncertainty and cognitive load on user trust in predictive decision making. In Human-Computer Interaction – INTERACT 2017 (pp. 23–39). Springer International Publishing.

Zorowitz, S., Rockhill, A. P., Ellard, K. K., Link, K. E., Herrington, T., Pizzagalli, D. A., Widge, A. S., Deckersbach, T., & Dougherty, D. D. (2019). The Neural Basis of Approach-Avoidance Conflict: A Model Based Analysis. eNeuro, 6(4). 10.1523/ENEURO.0115-19.2019

